# Prefrontal-amygdalar oscillations related to social interaction behavior in mice

**DOI:** 10.1101/2021.11.03.467135

**Authors:** Nahoko Kuga, Reimi Abe, Kotomi Takano, Yuji Ikegaya, Takuya Sasaki

## Abstract

The medial prefrontal cortex and amygdala are involved in the regulation of social behavior and associated with psychiatric diseases but their detailed neurophysiological mechanisms at a network level remain unclear. We recorded local field potentials (LFPs) from the dorsal medial PFC (dmPFC) and basolateral amygdala (BLA) while mice engaged on social behavior. We found that in wild-type mice, both the dmPFC and BLA increased 4–7 Hz oscillation power and decreased 30–60 Hz power when they needed to attend to another target mouse. In mouse models with reduced social interactions, dmPFC 4–7 Hz power further increased especially when they exhibited social avoidance behavior. In contrast, dmPFC and BLA decreased 4–7 Hz power when wild-type mice socially approached a target mouse. Frequency-specific optogenetic manipulations of replicating social approach-related LFP patterns restored social interaction behavior in socially deficient mice. These results demonstrate a neurophysiological substrate of the prefrontal cortex and amygdala related to social behavior and provide a unified pathophysiological understanding of neuronal population dynamics underlying social behavioral deficits.

## INTRODUCTION

The medial prefrontal cortex (mPFC) plays a central role in social behavior (Duncan & Owen, 2000; Wood & Grafman, 2003; Bicks *et al*., 2015) through functional interactions with the amygdala (Kumar *et al*., 2014; Adhikari *et al*., 2015; Bukalo *et al*., 2015; Tovote *et al*., 2015), a region that is reciprocally connected with the mPFC (Vertes, 2004; Hoover & Vertes, 2007; Adhikari *et al*., 2015) and plays central roles in emotional responses such as fear and anxiety. While a number of gene expression patterns and intracellular signaling pathways related to social behavior have been identified in the mPFC (Bicks *et al*., 2015; Schubert *et al*., 2015; Yan & Rein, 2021), a fundamental issue is how such molecular and cellular mechanisms are integrated to form organized mPFC and amygdalar neuronal population activity that cooperatively controls social behavior. Under pathological conditions, a number of studies have reported alterations in overall mPFC neuronal excitability and disruptions of mPFC-amygdala interactions in humans with psychiatric disorders with social behavior deficits such as autism spectrum disorders (ASD) and depression (Happe *et al*., 1996; Castelli *et al*., 2002; Pierce *et al*., 2004; Greicius *et al*., 2007; Drysdale *et al*., 2017) and animal models of these disorders (Brumback *et al*., 2018; Abe *et al*., 2019). It remains to be resolved which patterns of neuronal activity undergo pathological changes and cause social behavior deficits.

Neurophysiological signatures representing neuronal population activity are local field potential (LFP) signals, consisting of diverse oscillatory patterns that dynamically vary with attentional, motivational, and arousal states and entrain synchronous rhythmic spikes (Buzsaki, 2006). A recent study identified several patterns of prestress LFP power and coherence in frequency bands ranging from 1 to 50 Hz across whole brain regions differentiate stress vulnerability and susceptibility (Hultman *et al*., 2018). Especially, in the prefrontal-amygdalar circuit, theta-range (4–12 Hz) LFP signals are dominant in the expression of emotional behavior (Caliskan & Stork, 2019) such as fear retrieval (Seidenbecher *et al*., 2003; Likhtik *et al*., 2014; Stujenske *et al*., 2014; Dejean *et al*., 2016; Karalis *et al*., 2016; Ozawa *et al*., 2020) and anxiety (Adhikari *et al*., 2010; Likhtik *et al*., 2014). In addition, mPFC LFP power at a similar frequency (2–7-Hz) band influences oscillatory activity in the amygdala and ventral tegmental area during stress experiences (Hultman *et al*., 2016) and predicts vulnerability to mental stress in individual animals (Kumar *et al*., 2014). While these studies imply that social behavior is mediated by oscillatory signals at relatively low frequency (up to 10 Hz) bands in the prefrontal-amygdalar circuit, their causal relationship and pathological changes remain fully elusive. Addressing these issues is critical for a unified understanding of neurophysiological mechanisms at a neuronal network level underlying social behavior and its deficits.

Here, we analyzed changes in LFP signals from the dorsal mPFC (dmPFC) and basolateral amygdala (BLA) among wild-type mice and mouse models with social behavioral deficits in a social interaction (SI) test. By extracting detailed animal’s behavioral patterns on a moment-to-moment basis that potentially reflect increased and decreased motivation for social interactions, we discovered prominent changes in dmPFC-BLA LFP signals that specifically varied with social behavior. Optogenetic experiments verified a causal relationship between these oscillatory signals and social interaction behavior, highlighting the importance of frequency-specific manipulations of neuronal activity in the dmPFC-BLA circuit.

## RESULTS

### Changes in dmPFC and BLA LFP power in a social interaction test

C57Bl/6J mice were tested in a conventional SI test in which they freely interacted with an empty cage and a cage containing a target CD-1 mouse for 150 s, termed a no target and a target session, respectively (Fig. 1A). The degree of social interactions for each mouse was quantified as a social interaction (SI) ratio, which refers to the ratio of stay duration in an interaction zone (IZ) in a target session to that in a no target session. Consistent with previous observations (Golden *et al*., 2011; Venzala *et al*., 2012; Ramaker & Dulawa, 2017), the majority of wild-type mice (11 out of 14) exhibited SI ratios of more than 1 (Fig. 1B), demonstrating their motivation for social interactions. From the mice performing the SI tests, LFP signals were simultaneously recorded from the dmPFC, corresponding to the prelimbic (PL) region, and the BLA using an electrode assembly (Fig. 1C and 1D and Supplementary Fig. 1). The locations of individual electrodes were confirmed by a postmortem histological analysis. To compute an overall tendency of LFP power changes, a Fourier transformation analysis was applied to LFP signals from each entire session. We found a significant increase and decrease in dmPFC power at 4–7 Hz and 30–60 Hz, respectively, in a target session compared with a no target session (Fig. 1E; *n* = 14 mice, paired *t*-test between the target and no target sessions, *q* < 0.05, FDR corrected). For detailed quantification of power changes across mice, LFP power at each frequency band in each region was z-scored in each mouse. Overall, z-scored 4–7 Hz and 30–60 Hz power in both the dmPFC and BLA during the target session was significantly higher and lower, respectively, than that during the no target session (Fig. 1F, dmPFC: *n* = 14 mice, 4– 7 Hz: *t*_13_ = 4.20, *P* = 0.0010; 30–60 Hz: *t*_13_ = 3.28, *P* = 0.0060; Fig. 1G, BLA: *n* = 6 mice, 4–7 Hz: *t*_5_ = 2.61, *P* = 0.047; 30–60 Hz: *t*_5_ = 3.06, *P* = 0.028, paired *t*-test; Fig. 2C, 2D, 3C, and 3D, indicated by $). The same analyses were applied to the same datasets at the other frequency bands, including 1–4 Hz, 7–10 Hz, 10–30 Hz, and 60–100 Hz bands, but no significant differences were found between the target and no target sessions (Supplementary Fig. 2A and 2B). These results suggest that dmPFC-BLA 4–7 Hz and 30– 60 Hz power specifically become higher and lower, respectively, when mice are exposed to an environment with the other target mouse.

**Figure 1.**
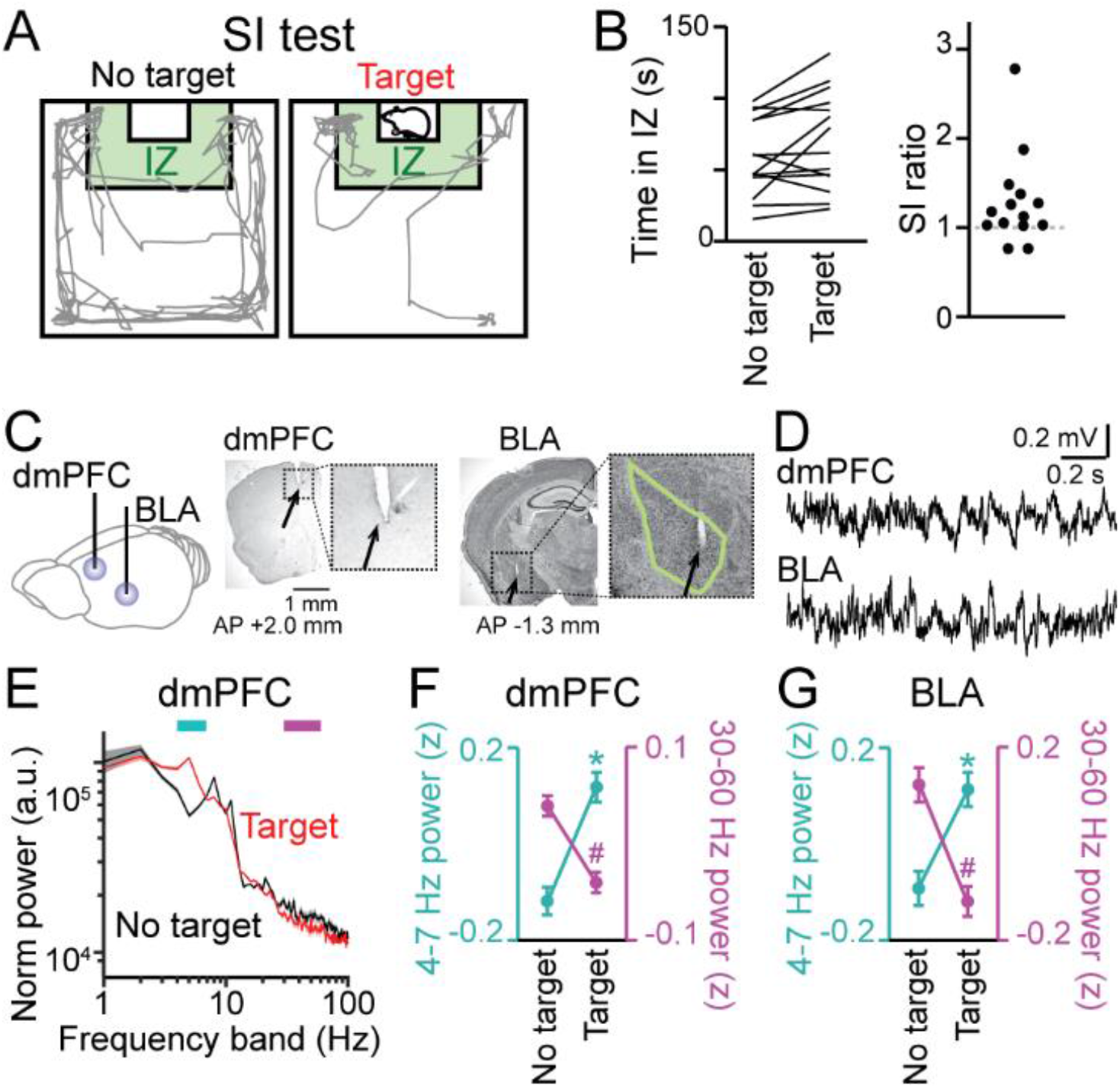
Changes in dorsal medial prefrontal cortex (dmPFC) and basolateral amygdala (BLA) local field potential (LFP) signals in a social interaction (SI) session. (A) SI test with an interaction zone (IZ; labeled in green). Movement trajectories (gray lines) of a wild-type mouse are superimposed. (B) (Left) Occupancy time in the IZ. Each line indicates an individual mouse (*n* = 14 wild-type mice). (Right) SI ratios computed from the occupancy time. Each dot represents an individual mouse. (C) (Left) LFPs were recorded from the dmPFC and BLA. (Right) Histological confirmation of electrode locations (arrows). The dotted boxes are magnified in the right panels. The green line shows the contour of the BLA. The details of electrode locations are shown in Supplementary Figure 1. (D) Typical LFP signals from the dmPFC and BLA. (E) Comparison of dmPFC LFP power spectrograms between target (red) and no target (black) sessions in a mouse. Data are presented as the mean ± SEM. Cyan and magenta bars above represent 4–7 Hz and 30–60 Hz bands, respectively. (F) Comparison of dmPFC 4–7 Hz (cyan) and 30–60 Hz (magenta) power between the target and no target sessions (*n* = 14 mice). Data are presented as the mean ± SEM. * and # represent a significant increase and decrease in the target session, respectively (*P* < 0.05, paired *t*-test vs no target). (G) Same as F but for the BLA (*n* = 6 mice).

**Figure 2.**
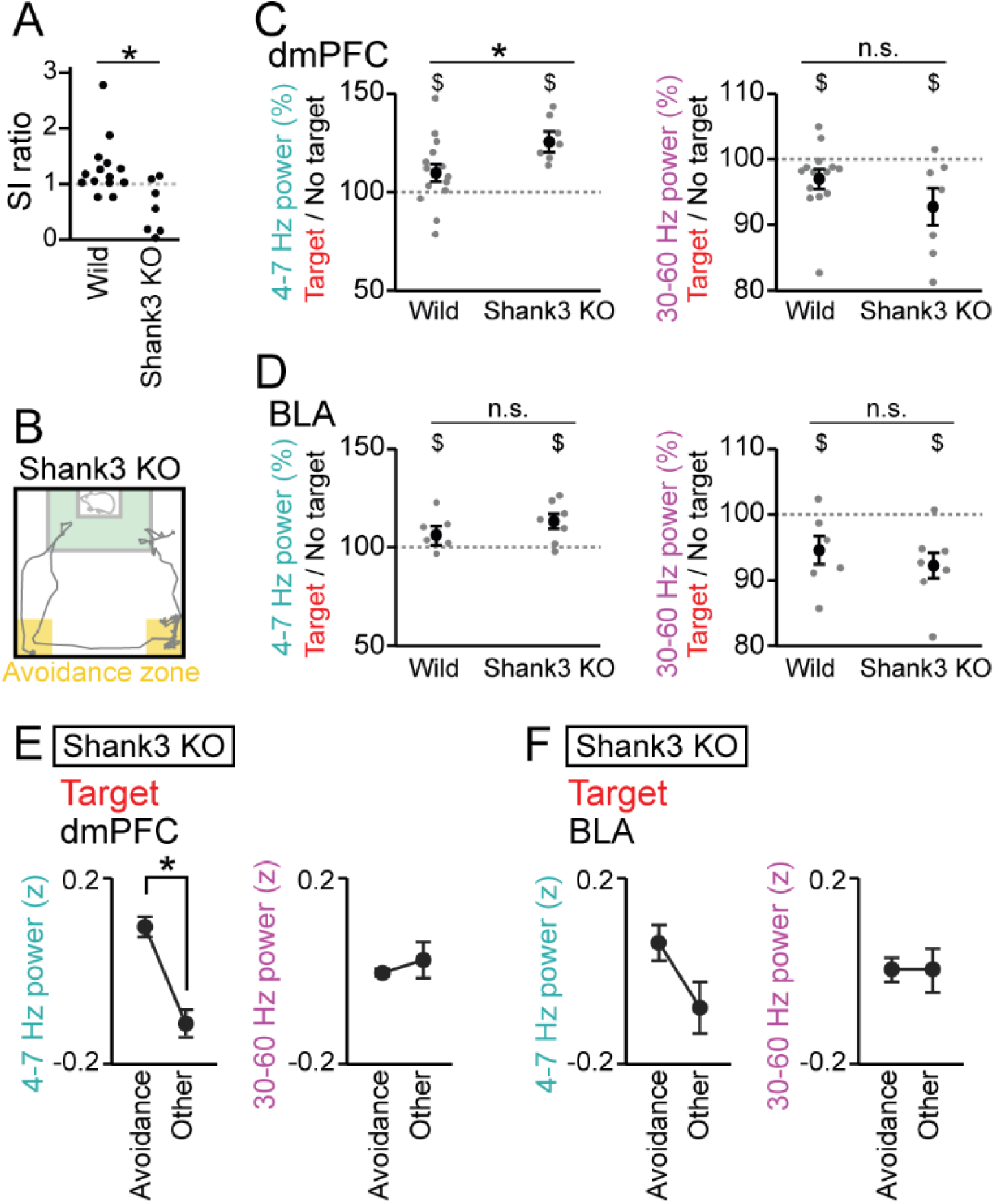
Increases in dmPFC 4–7 Hz power during social avoidance in Shank3 KO mice. (A) SI ratios for Shank3 KO mice. Each dot represents an individual animal. The data from wild-type mice similar to those shown in Figure 1B are presented for comparison. \**P* < 0.05 versus wild, Mann-Whitney U test. (B) Movement trajectory of a Shank3 KO mouse in a target session. The orange areas represent the avoidance zones. (C) The percentages of dmPFC 4–7 Hz (left) and 30–60 Hz (right) LFP power in the target session relative to those in the no target session (*n* = 14 wild and 7 Shank3 KO mice). Data are presented as the mean ± SEM. Each gray dot represents an individual data points. ^$^*P* < 0.05, no target vs target within a group. \**P* < 0.05, Mann-Whitney U test across groups. (D) Same as C but for the BLA (*n* = 6 and 7 mice). (E) Comparisons of dmPFC 4–7 Hz (left) and 30–60 Hz (right) power between the avoidance zones and the other areas in Shank 3 KO mice (*n* = 7 Shank3 KO mice). Data are presented as the mean ± SEM. \**P* < 0.05, paired *t*-test. (F) Same as F but for the BLA (*n* = 7 Shank3 KO mice).

**Figure 3.**
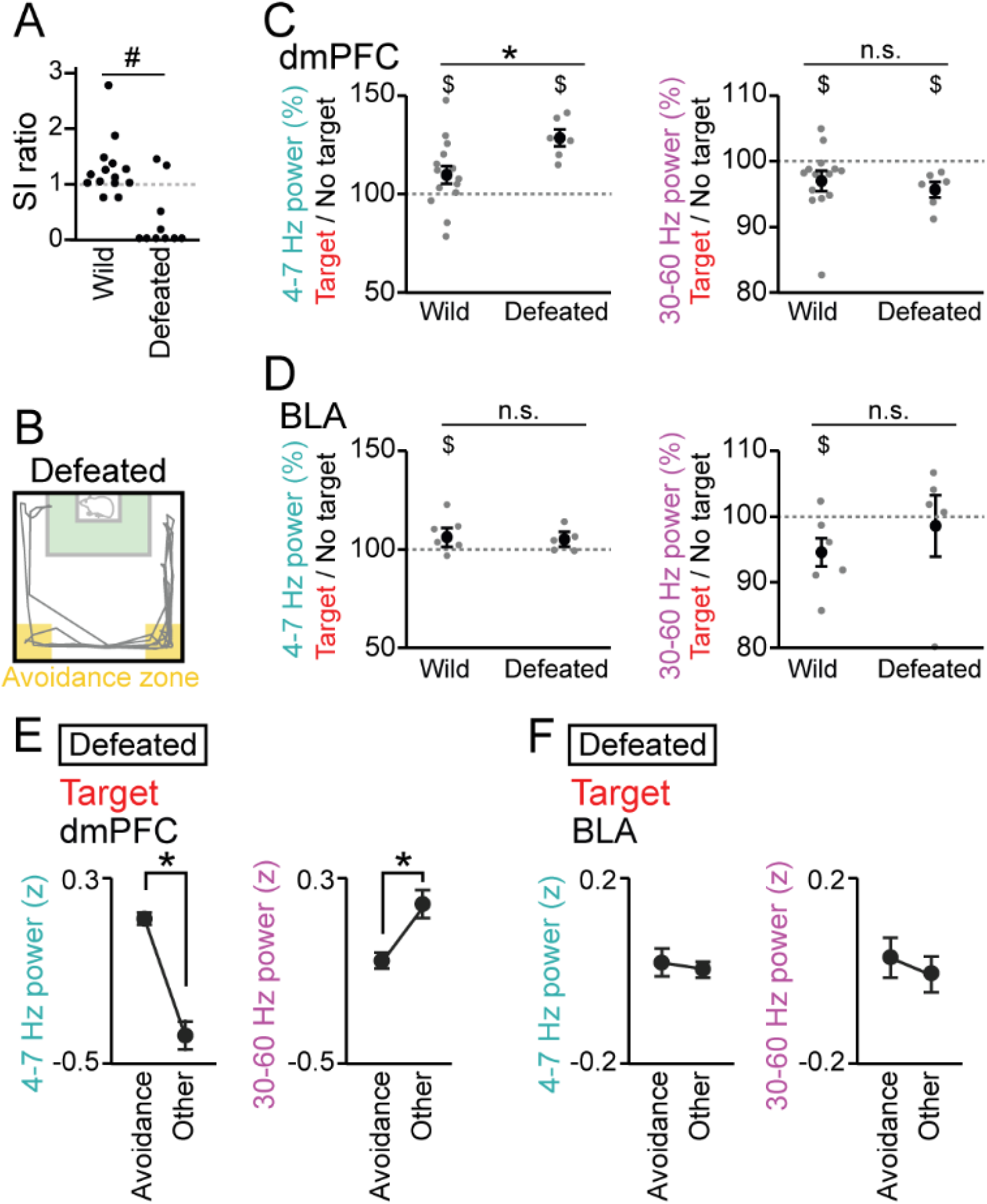
Increases in dmPFC 4–7 Hz power during social avoidance in defeated mice. (A) SI ratios for defeated mice. \**P* < 0.05 versus wild, Mann-Whitney U test. (B) A movement trajectory of a defeated mouse in a target session. (C) The percentages of dmPFC 4–7 Hz (left) and 30– 60 Hz (right) LFP power in the target session relative to those in the no target session (*n* = 14 wild and 6 defeated mice). Data are presented as the mean ± SEM. Each gray dot represents an individual data point. ^$^*P* < 0.05, no target vs target within a group. \**P* < 0.05, Mann-Whitney U test across groups. (D) Same as C but for the BLA (*n* = 6 wild and 5 defeated mice). (E) Comparisons of dmPFC 4–7 Hz (left) and 30–60 Hz (right) power between the avoidance zones and the other areas in defeated mice (*n* = 6 defeated mice). Data are presented as the mean ± SEM. \**P* < 0.05, paired *t*-test. (F) Same as F but for the BLA (*n* = 5 defeated mice).

### Increases in dmPFC 4–7 Hz power during social avoidance in socially deficient mouse models

We then examined whether these LFP signals are altered in mice with reduced social interaction. The same tests were performed on Shank3 null mutant mice, termed Shank3 knockout (KO) mice, which have been reported to exhibit repetitive grooming behavior and social interaction deficits, mimicking symptoms associated with ASD (Peca *et al*., 2011; Mei *et al*., 2016). The SI ratios in 7 Shank3 KO mice were significantly lower than those in the 14 wild-type mice (Fig. 2A and 2B, *Z* = 2.65, *P* = 0.016, Mann-Whitney U test followed by Bonferroni correction). We recorded LFP signals from the dmPFC and BLA of these Shank3 KO mice and found significant increases and decreases in dmPFC LFP power during the target session at the 4–7 Hz and 30–60 Hz bands, respectively, similar to the wild-type mice (Fig. 2C, dmPFC: *n* = 7 mice, 4–7 Hz, *t*_6_ = 6.04, *P* = 9.2 × 10^-4^; 30–60 Hz; *t*_6_ = 2.62, *P* = 0.039; Fig. 2D, BLA: *n* = 7 mice, 4–7 Hz, *t*_6_ = 2.93, *P* = 0.026; 30–60 Hz; *t*_6_ = 3.52, *P* = 0.013, paired *t*-test, indicated by $). Moreover, the dmPFC 4–7 Hz increases during the target session in the Shank3 KO mice were significantly larger than those observed in the wild-type mice (Fig. 2C, *Z* = 2.20, *P* = 0.028, Mann-Whitney U test), whereas no differences were observed for the changes in dmPFC 30–60 Hz power (*Z* = 1.23, *P* = 0.22, Mann-Whitney U test) and BLA power (Fig. 2D, 4–7 Hz, *P* = 0.45; 30–60 Hz, *P* = 0.73, Mann-Whitney U test). These results suggest that the increases in dmPFC 4– 7 Hz power during a target session are more prominent in Shank3 KO mice with decreased social behavior.

During the target session, the Shank3 KO mice spent substantial (24.7 ± 6.7%) time near the corners of the field opposing the target mice, termed avoidance zones (Fig. 2B), which are considered as social avoidance behavior that potentially represents decreased motivation for social behavior (Golden *et al*., 2011). To examine whether such typical social avoidance behavior is associated with LFP patterns at the frequency bands found above, we extracted 4–7 Hz and 30–60 Hz LFP power in the avoidance zones in the target session. DmPFC 4–7 Hz was significantly increased when the Shank3 KO mice stayed within the avoidance zone, compared with the other areas (Fig. 2E, dmPFC: *n* = 7 mice, 4– 7 Hz, *t*_5_ = 4.39, *P* = 0.0071; 30–60 Hz; *t*_5_ = 0.33, *P* = 0.75; Fig. 2F, BLA: *n* = 7 mice, 4–7 Hz, *t*_5_ = 1.62, *P* = 0.16; 30–60 Hz; *t*_5_ = 0.17, *P* = 0.87, paired *t*-test). These results demonstrate that a 4–7 Hz power increase occurs in the dmPFC when Shank3 KO mice exhibit avoidance behavior.

We next tested whether socially defeated mice with reduced social interaction consecutive days, termed defeated mice. SI ratios of the 6 defeated mice were significantly lower than those in the 14 wild-type mice (Fig. 3A and 3B; *Z* = 4.10, *P* = 8.1 × 10^-5^, Mann-Whitney U test followed by Bonferroni correction). Similar to the wild-type and Shank3 KO mice, these defeated mice exhibited a significantly larger increase in dmPFC 4–7 Hz power during the target session than during the no target session (Fig. 3C, dmPFC: *n* = 6 mice, 4–7 Hz, *t*_5_ = 6.93, *P* = 9.6 × 10^-4^; 30–60 Hz; *t*_5_ = 3.77, *P* = 0.013; Fig. 3D, BLA: *n* = 5 mice, 4–7 Hz: *t*_4_ = 1.88, *P* = 0.14; 30–60 Hz: *t*_4_ = 0.33, *P* = 0.76, paired *t*-test, indicated by $). In addition, similar to the Shank3 KO mice, the dmPFC 4–7 Hz increase in the defeated mice was significantly larger than that observed from the wild-type mice (Fig. 3C, dmPFC: 4–7 Hz: *Z* = 2.27, *P* = 0.023; 30–60 Hz: *Z* = 1.36, *P* = 0.17; Fig. 3D, BLA: 4–7 Hz: *P* = 0.79; 30–60 Hz: *P* = 0.25, Mann-Whitney U test) and dmPFC 4–7 Hz power in the defeated mice was significantly increased in the avoidance zone (Fig. 3E, dmPFC: *n* = 6 mice, 4–7 Hz, *t*_5_ = 11.68, *P* = 8.8 × 10^-5^; 30–60 Hz; *t*_5_ = 3.45, *P* = 0.018; Fig. 3F, BLA: *n* = 5 mice, 4–7 Hz, *t*_4_ = 0.47, *P* = 0.66; 30–60 Hz; *t*_4_ = 0.46, *P* = 0.67, paired *t*-test). These results demonstrate that a dmPFC 4–7 Hz power increase during social avoidance also occurs in depression model mice. Taken together, our results from the two mouse models suggest that dmPFC 4–7 Hz power increases during social avoidance behavior are a common hallmark across socially deficient mouse models, contributing to their overall increase in dmPFC 4–7 Hz power during a target session.

### Changes in dmPFC LFP power during social approach behavior

The social avoidance-related increases in dmPFC 4–7 Hz power implied that social interaction behavior may be associated with dmPFC 4–7 Hz power changes. To test this possibility, we first compared LFP power between when the wild-type normal mice stayed within and were outside the IZ as conventional measures in an SI test. However, no significant changes in 4–7 Hz and 30–60 Hz power in the dmPFC and BLA during the target session were detected between the IZ and the other areas (Supplementary Fig. 3A and 3B; dmPFC, *n* = 14 mice; 4–7 Hz: *t*_13_ = 0.76, *P* = 0.46; 30–60 Hz: *t*_13_ = 1.38, *P* = 0.19; BLA, *n* = 6 mice; 4–7 Hz: *t*_5_ = 0.76, *P* = 0.48; 30–60 Hz: *t*_5_ = 0.02, *P* = 0.98). While this analysis focused on the entire period during which the mice stayed in the IZ, their behavioral patterns within the IZ were not consistent across time; mice actively approach or interact with a target mouse in some periods, reflecting high motivation, whereas they occasionally turn around, move away from a target mouse, or continue to stay at a location in an IZ, possibly reflecting no strong motivation for social interaction. These behavioral observations indicate that animals’ motivation toward and salience regarding the other mouse are not equivalent even when they are similarly located in an IZ.

The results suggest that changes in dmPFC-BLA oscillations are not simply explained by where the mice stayed in the SI test. We further analyzed how LFP patterns are associated with their instantaneous behavior defined every 1 s. First, running and stop periods were defined when animal’s moving speed was more than 5 cm/s and less than 1 cm/s, respectively. Comparisons of distributions of LFP power between the two periods demonstrated that 4–7 Hz power during running periods was significantly lower than that during stop periods both in the dmPFC and BLA (Supplementary Fig. 3C; dmPFC, *n* = 14 mice, *t*_13_ = 0.76, *P* = 0.46; BLA, *n* = 6 mice, *t*_5_ = 0.76, *P* = 0.48), whereas 30–60 Hz power exhibited opposite changes (Supplementary Fig. 3C; dmPFC, *n* = 14 mice, *t*_13_ = 1.38, *P* = 0.19; BLA, *n* = 6 mice, *t*_5_ = 0.02, *P* = 0.98), suggesting that locomotion is a crucial factor to affect these LFP power changes. Next, to more precisely define behavioral patterns related to increased motivation for social interaction, we defined social approach behavior as the time during which the mice approached the cage (within the half of the field containing the cage) with their cage-oriented moving directions θ less than 90° in the target session (Fig. 4A–C). Assuming that mice potentially exhibited the highest motivation and salience during an initial bout of a social approach, this definition was restricted to the initial 5-s periods of social approach behavior. As a control behavior against approach behavior, we defined leaving behavior as the time during which the mice left from the cage with their cage-oriented moving directions θ more than 90° in the target session. No significant differences in the distributions of moving speed were found between the approach behavior and leaving behavior (Fig. 4D; *Z* = 0.85, *P* = 0.39, Mann-Whitney U test). These results confirm that approach and leaving behavior is not explained by speed or locomotion itself, allowing us to compare LFP power between the two behavioral periods without being affected by moving speed. Both in the dmPFC and BLA, 4–7 Hz power was significantly decreased during the approach behavior compared with the leaving behavior (Fig. 4F; dmPFC: 4–7 Hz: *t*_13_ = 2.71, *P* = 0.018; 30–60 Hz: *t*_13_ = 0.47, *P* = 0.65; Fig. 4G; BLA: 4–7 Hz: *t*_5_ = 2.83, *P* = 0.037; 30–60 Hz: *t*_5_ = 1.01, *P* = 0.36, paired *t*-test). No significant differences in LFP power at the other frequency bands (1–4 Hz, 7–10 Hz, 10–30 Hz, and 60–100 Hz bands) were observed between the approach and leaving behavior (Supplementary Fig. 2C and 2D). The reductions of dmPFC 4–7 Hz power during social interaction behavior are consistent with the opposite changes (the increases in dmPFC 4–7 Hz power) observed during social avoidance behavior in the socially deficient mice (Fig. 2E and 3E). Taken together, our results suggest that changes in dmPFC 4–7 Hz oscillations are a key neuronal substrate to bidirectionally modulate social interaction behavior.

**Figure 4.**
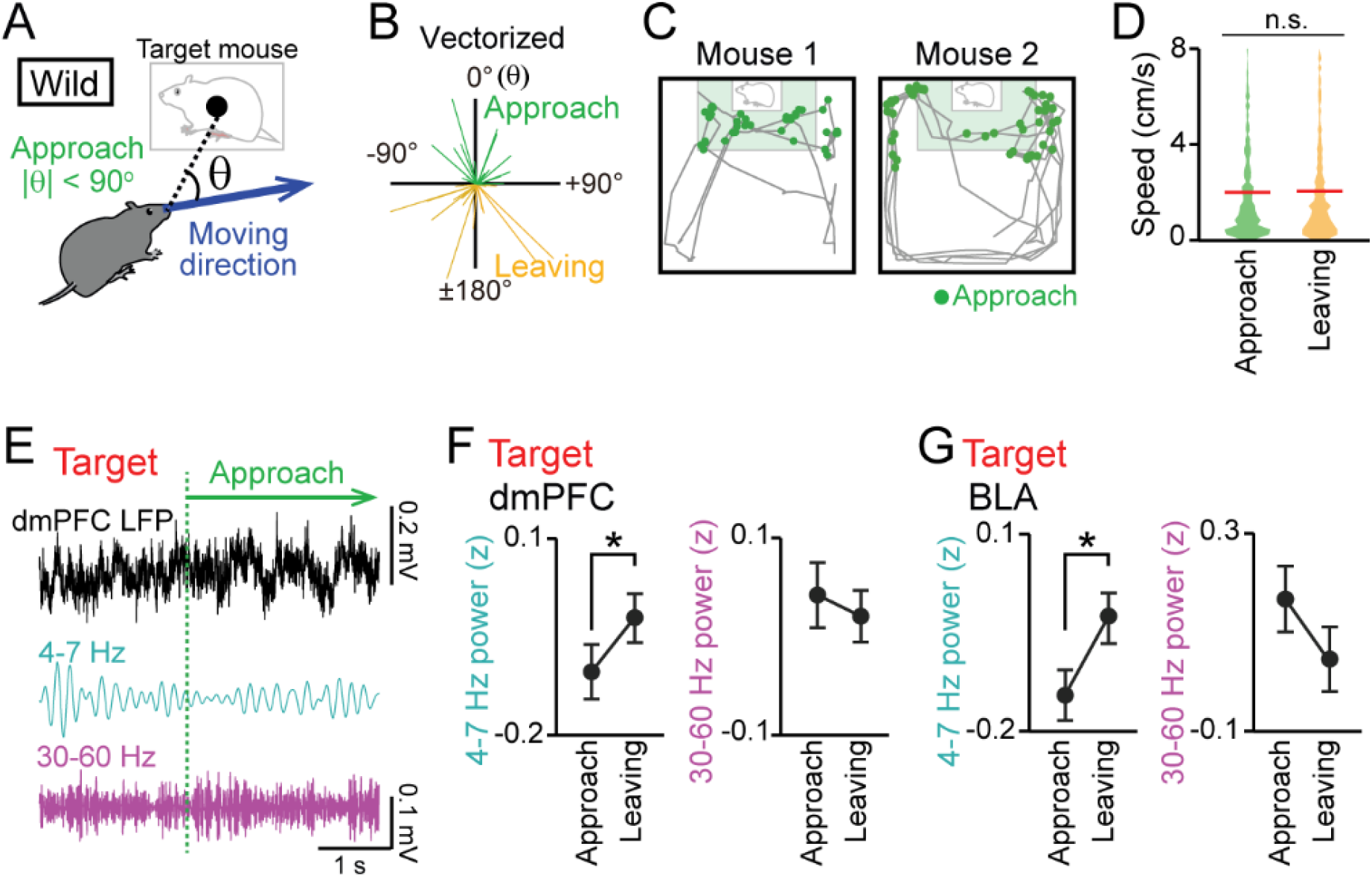
Decreases in dmPFC 4–7 Hz LFP power during social approach behavior. (A) Social approach and leaving behavior in wild-type mice was defined when absolute cage-oriented moving directions (|θ|) were less than and more than 90°, respectively, in the half of the field containing the IZ. (B) A polar plot of vectorized instantaneous animal trajectories (bin = 1 s) as a function of cage-oriented moving direction. (C) Trajectories from two representative mice (gray). Green dots represent social approach behavior. (D) Distributions of moving speed during approach and leaving behavior (*n* = 644 and 577). The red lines show the average. *P* > 0.05, Mann-Whitney U test. (E) Unfiltered and bandpass (4–7 Hz and 30–60 Hz)-filtered dmPFC LFP traces in the target session. The green line indicates the onset of a social approach behavior. (F) Comparison of dmPFC 4–7 Hz (left) and 30–60 Hz (right) power between approach and leaving behavior (*n* = 14 mice). Data are presented as the mean ± SEM. \**P* < 0.05, paired *t*-test. (G) Same as F but for the BLA (*n* = 6 mice).

### Neuronal spikes associated with LFP oscillations in the mPFC

We next analyzed how these LFP oscillations entrain spikes of individual mPFC neurons, defined by single-unit isolation. Following the criteria utilized in previous studies (Wilson *et al*., 1994; Tierney *et al*., 2004; Homayoun & Moghaddam, 2007), as shown in Figure 5A, putative regular-spiking (RS) excitatory pyramidal neurons were identified as neurons that had baseline spike rates lower than 10 Hz and spike widths longer than 0.6 ms (*n* = 48 neurons from 11 mice). On the other hand, putative fast-spiking (FS) interneurons were identified as neurons that had baseline spike rates higher than 10 Hz (*n* = 9 neurons from 6 mice). While these criteria might define a minority of interneurons as putative RS neurons, it was unlikely that true pyramidal neurons are misclassified as putative FS neurons (Wilson *et al*., 1994; Tierney *et al*., 2004; Homayoun & Moghaddam, 2007). We tested whether these mPFC neurons exhibited spike patterns phase-locked to the 4–7 Hz oscillations (Fig. 5B). All spike analyses were restricted to a target session. An example putative FS neuron shown in Figure 5B exhibited apparent spike rate changes corresponding to altering phases in the 4–7 Hz oscillations. For each neuron, the degree of spike phase locking was quantified by computing the mean vector length (MVL). In the example neuron, the MVL was 0.12. To assess the significance of each MVL, we created shuffled datasets in which spike timing was randomized within the session and MVL was similarly computed from 1000 shuffled datasets, termed MVL_shuffled_. The MVL of an original data was considered to be significant (*P* < 0.05) when the MVL was higher than the top 95% of the corresponding MVL_shuffled_. According to this criterion, the MVL of the example neuron was higher than the corresponding 1000 MVL_shuffled_ (*P* < 0.001), demonstrating that these neuronal spikes were entrained by the 4–7 Hz oscillations. For each neuron, the MVL was compared between approach and leaving behavior (Fig. 5C). Of the 48 and 9 putative RS and FS neurons tested, 5 (10.4%) and 3 (33.3%) neurons showed significant MVL in both approach and leaving behavior (the neurons indicated by the red lines in Figure 5C). These neurons were considered to show phase locking spikes irrespective of behavior and excluded from further statistical analyses. Overall, FS neurons showed significantly higher MVL for the 4–7 Hz oscillations during leaving periods, compared with approach periods (*n* = 6 neurons; *t*_5_ = 3.15, *P* = 0.025, paired *t*-test), whereas RS neurons did not show such significance (*n* = 43 neurons; *t*_42_ = 1.69, *P* = 0.69). These results demonstrate that FS neurons alter their entrainment to the 4–7 Hz oscillations depending on animal’s behavioral patterns. The same analyses were applied to the other frequency bands (1–4 Hz, 4–7 Hz, 7–15 Hz, 15–30 Hz, and 30–60 Hz) but no significant differences were observed between approach and leaving behavioral periods (Supplementary Fig. 4; *P* > 0.05, paired *t*-test). Taken together, these results suggest that mPFC PS neurons are more preferentially entrained by the 4–7 Hz oscillations during leaving behavior than approach behavior. In other words, the entrainment of mPFC neurons to the 4–7 Hz oscillations is disrupted when mice engage in social approach behavior.

**Figure 5.**
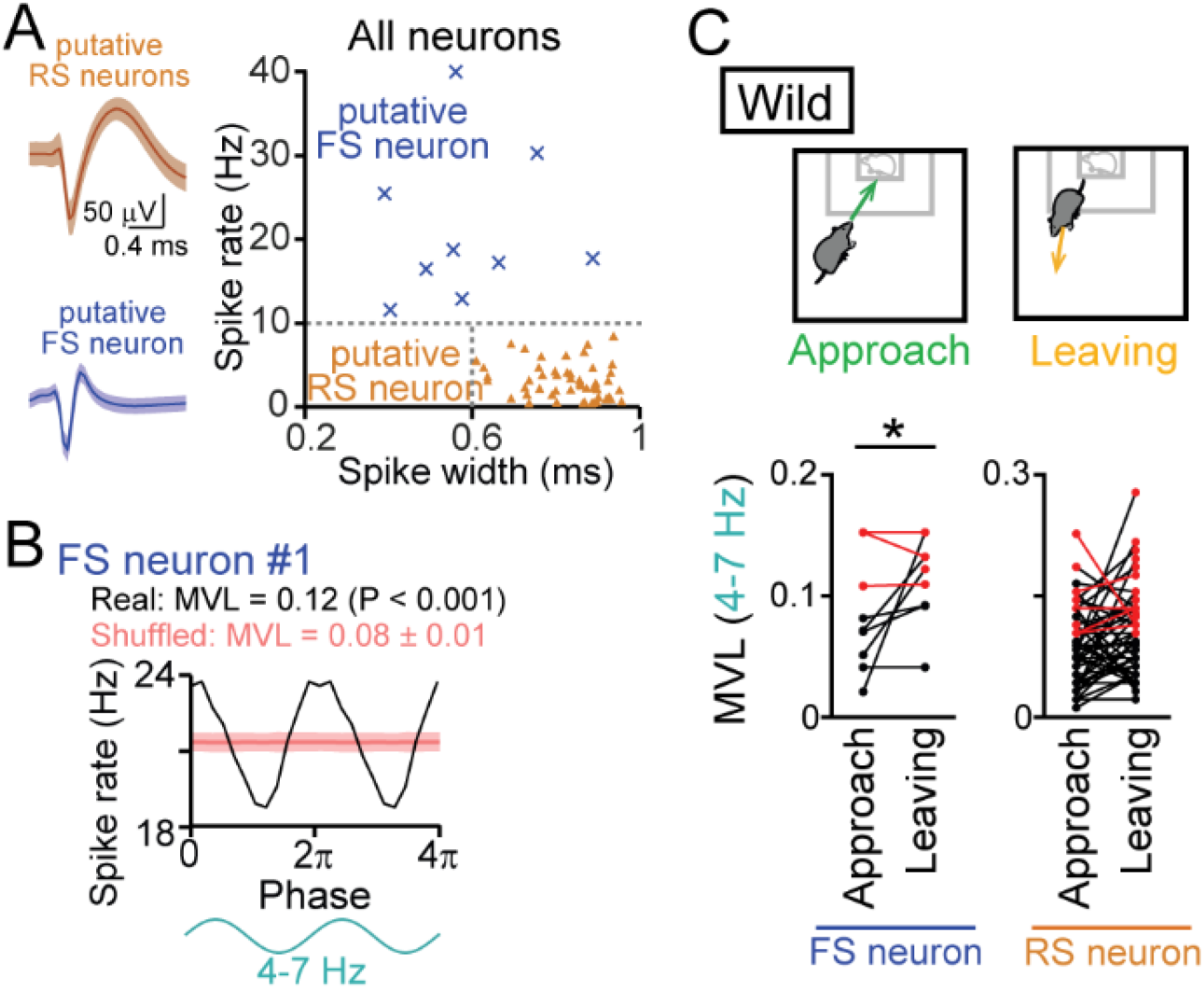
Entrainment of oscillatory spike patterns of mPFC neurons. (A) (Left) Typical spike waveforms of a putative regular-spiking (RS) neuron (top) and a putative fast-spiking (FS) neuron (bottom). Data are presented as the mean ± SD. (Right) For individual neurons in wild-type mice, baseline spike rates and spike width are plotted. Each dot represents an individual neuron. Neurons plotted in orange and cyan regions are classified as putative excitatory RS pyramidal neurons (triangle, *n* = 48 neurons) and inhibitory FS interneurons (cross-mark, *n* = 9 neurons), respectively. (B) A representative putative FS neuron that fired time-locked to the 4–7-Hz LFP oscillations. Instantaneous spike rates of this neuron are plotted against the phase of 4–7-Hz oscillatory cycles. From a phase-spike distribution, a mean vector length (MVL) was computed as 0.12. The red line and the shaded area represent the mean and SD computed from the corresponding 1000 shuffled datasets. (C) Comparisons of MVL for the 4–7-Hz LFP oscillations between approach and leaving behavior. RS and FS neurons were separately analyzed. Each dot and line represent each neuron. The red dots indicate significant MVL, computed from shuffled datasets. The red lines indicate neurons showing significant MVL in both of the periods, which were considered as behavior-irrelevant phase-locked neurons and excluded from the statistical analyses. *P* < 0.05, paired *t*-test for the datasets shown in black.

### Restoration of social interaction by optogenetic manipulation of dmPFC 4–7 Hz power

The observations that dmPFC 4–7 Hz power was reduced during social approach behavior imply that replicating such dmPFC LFP patterns may potentially facilitate social interaction behavior. To address this idea, a technique to reduce dmPFC 4–7 Hz oscillations is needed. Based on our observations that the entrainment of mPFC inhibitory neuronal spikes to the 4–7 Hz oscillations dynamically varies with social behavior, we sought to develop a method to alter dmPFC 4–7 Hz oscillations by manipulating dmPFC inhibitory neurons. Here, we focused on parvalbumin (PV)-positive interneurons, a major type of interneurons as this cell type has been reported to be crucial for the generation of cortical gamma-range (30–60 Hz) oscillations (Whittington *et al*., 1995; Bartos *et al*., 2007; Cardin *et al*., 2009; Sohal *et al*., 2009; Buzsaki & Wang, 2012; Nakamura *et al*., 2015). To selectively control the activity of PV-positive interneurons, we utilized optogenetic tools with PV-Cre mice. While we noted that not all interneurons in the dmPFC express PV, we took this optogenetic approach as a means to potentially alter dmPFC neuronal oscillations at these frequency bands. The PV-Cre mice were first subjected to chronic social defeat stress and then injected with a Cre-inducible viral construct, AAV5-EF1a-DIO-ChR2-eYFP or AAV5-EF1a-DIO-eYFP, into the dmPFC so that PV interneurons selectively expressed ChR2-YFP or YFP, respectively (Fig. 6A). In addition, an optic cannula and recording electrodes were implanted into the identical region of the dmPFC, and additional recording electrodes were implanted into the BLA. We first sought to identify appropriate photostimulation protocols by simultaneous LFP recordings of the dmPFC and BLA. Two weeks after surgery, photostimulation at 4, 10, and 40 Hz was applied in defeated mice expressing ChR2-YFP (Fig. 6B), a frequency corresponding to 4–7 Hz, an intermediate frequency, and a frequency corresponding to 30–60 Hz, respectively. The width of each photostimulation pulse was set to half the pulse intervals: 125, 50, and 12.5 ms for 4, 10, and 40 Hz, respectively, meaning that the total duration of applied photostimulation was equivalent in all protocols. Photostimulation at 4 Hz induced a significant 19.5 ± 9.0% power increase in the corresponding frequency (4–7 Hz) band in the dmPFC (Fig. 6B and 6C; 4–7 Hz: *t*_15_ = 2.44, *P* = 0.045; 30–60 Hz: *t*_15_ = 2.18, *P* = 0.066, paired *t*-test), consistent with the entrainment of inhibitory neuronal spikes to 4–7 Hz oscillations. Photostimulation at 10 Hz induced a small (7.2 ± 2.2%) but significant power increase at the 30–60 Hz band in the dmPFC (4–7 Hz: *t*_15_ = 1.40, *P* = 0.20; 30–60 Hz: *t*_15_ = 3.28, *P* = 0.014). Photostimulation at 40 Hz induced significant 24.8 ± 14.1% and 2.8 ± 0.9% power increases at the corresponding frequency (30–60 Hz) band in the dmPFC and BLA, respectively (Fig. 6B and 6C, dmPFC: *t*_15_ = 2.91, *P* = 0.023; Fig. 6D, BLA: *t*_15_ = 3.27, *P* = 0.017). In addition, the 40-Hz photostimulation induced significant 17.5 ± 5.4% and 6.4 ± 2.6% power decreases in the 4–7 Hz band in the dmPFC and BLA, respectively (Fig. 6B and 6C, dmPFC: *t*_15_ = 3.24, *P* = 0.014; Fig. 6D, BLA: *t*_15_ = 2.49, *P* = 0.047). The bidirectional changes in 4–7 Hz and 30–60 Hz power in the dmPFC-BLA circuit at 40-Hz photostimulation are consistent with those observed during social approach behavior, as shown in Figure 4, suggesting that it was possible to replicate such social interaction-related neuronal activity.

**Figure 6.**
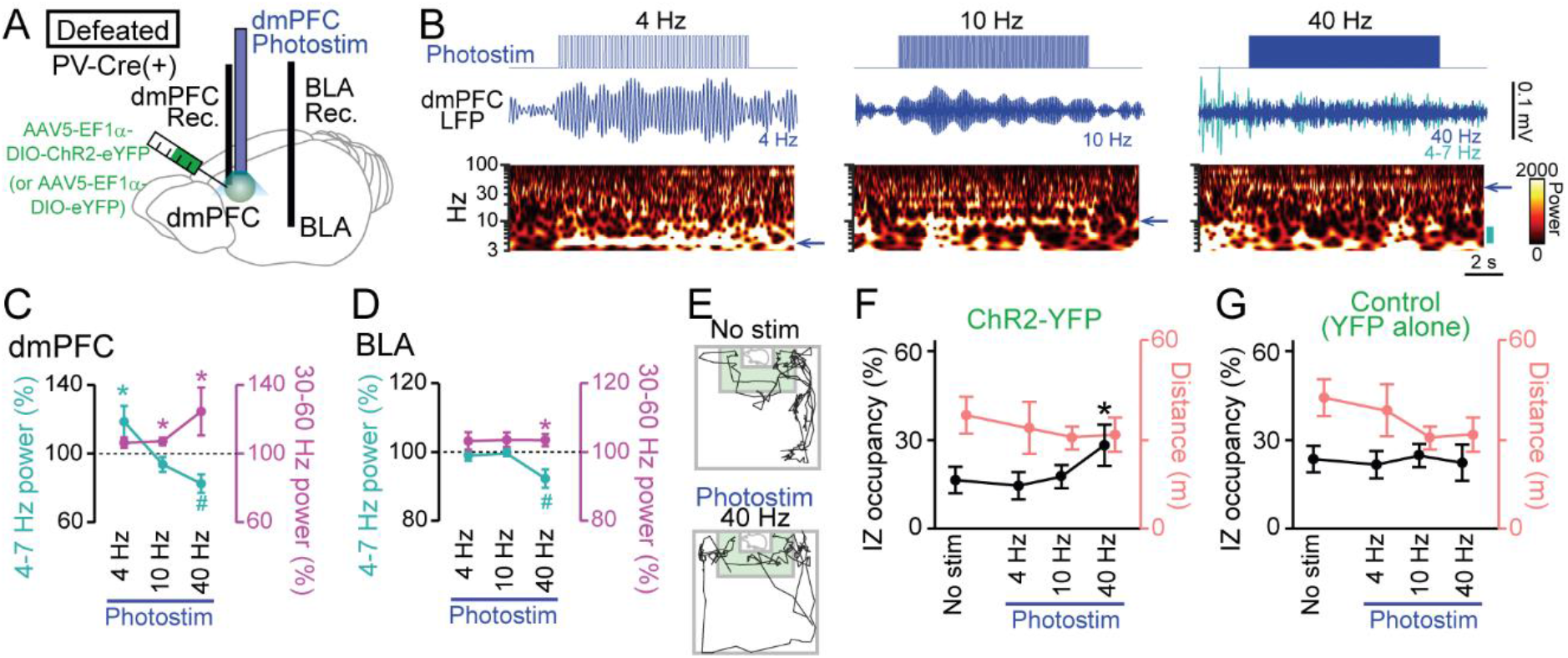
Restoration of social approach behavior by optogenetic photostimulation with a decrease in 4–7 Hz power and an increase in 30–60 Hz power in the dmPFC. (A) Schematic illustration. PV-positive interneurons expressing ChR2-YFP (or YFP alone) in the dmPFC were optogenetically stimulated, while LFP signals were recorded from the dmPFC and BLA in PV-Cre mice subjected to social defeat stress. (B) (Top) Photostimulation at 4, 10, and 40 Hz for 10 s in defeated mice injected with AAV5-EF1a-DIO-ChR2-eYFP. (Middle and bottom) Representative dmPFC LFP traces filtered at the frequency band similar to that of photostimulation (blue arrows) and its wavelet spectrum. At 40-Hz photostimulation, the LFP trace filtered at 4–7 Hz is superimposed as the cyan trace. The cyan bar beside the wavelet spectrum represents the 4–7 Hz band. (C) Average dmPFC 4–7 Hz (cyan) and 30–60 Hz (magenta) power changes by dmPFC photostimulation in defeated mice injected with AAV5-EF1a-DIO-ChR2-eYFP. (*n* = 7 mice). * and # represent a significant increase and decrease, respectively (*P* < 0.05, paired *t*-test versus baseline). (D) Same as C but for BLA power. (E) Movement trajectories of a defeated mouse in a target session with no stimulation and 40-Hz photostimulation. (F) The percentage of time spent in the IZ (left axis, black) and total travel distance (right axis, thin red) in defeated mice injected with AAV5-EF1a-DIO-ChR2-eYFP (*n* = 7 mice). Data are presented as the mean ± SEM. \**P* < 0.05, paired *t*-test followed by Bonferroni correction. (G) Same as F but for defeated mice injected with AAV5-EF1a-DIO-eYFP (*n* = 5 mice).

The defeated PV-Cre mice were tested in a target session with these photostimulation protocols at each frequency band (Fig. 6E). A test day includes a sequence of a target session with no photostimulation (no stim), and target sessions with 4-Hz, 10-Hz, and 40-Hz photostimulation. The order of sessions was similar in all mice. In all the sessions, photostimulation was applied for 2 s every 30 s and overall changes in the animal’s social interactions throughout the session were examined. In the mice expressing ChR2-YFP, 40-Hz photostimulation significantly increased occupancy time in the IZ, compared with no photostimulation (*n* = 7 mice; *t*_6_ = 3.46, *P* = 0.040, paired *t*-test followed by Bonferroni correction), whereas no significant changes were observed by 4-Hz and 10-Hz photostimulation (4-Hz: *t*_6_ = 0.56, *P* > 0.99; 10-Hz: *t*_6_ = 0.25, *P* > 0.99, paired *t*-test followed by Bonferroni correction). There were no significant changes in total travel distance by these photostimulation conditions (Fig. 6E; *n* = 7 mice; *P* > 0.05 versus no stim, paired *t*-test followed by Bonferroni correction). On the other hand, in the mice expressing YFP alone, no significant effects were observed in any photostimulation conditions (*n* = 5 mice; 4-Hz: *t*_4_ = 2.05, *P* = 0.87; 10-Hz: *t*_4_ = 2.16, *P* = 0.83; 40-Hz: *t*_4_ = 2.51, *P* = 0.72, paired *t*-test followed by Bonferroni correction). These results from control experiments exclude the possibility that the increase in IZ occupancy by the 40-Hz photostimulation is simply due to an effect of elapsed time in the test condition. Overall, these results demonstrate that optogenetic inductions of a decrease in dmPFC-BLA 4–7 Hz power and an increase in 30–60 Hz power are sufficient to trigger social interaction behavior.

## DISCUSSION

In this study, we compared LFP signals from the dmPFC and BLA during an SI test among wild-type mice, Shank3 KO mice as a model of ASD, and socially defeated mice as a model of depression. Power spectrum analyses revealed that all mouse types tested exhibited prominent increases in dmPFC-BLA 4–7 Hz power and decreases in 30–60 Hz power throughout a target session compared with a no target session. The dmPFC 4–7 Hz power increase was prominent when socially deficient mouse models exhibited social avoidance behavior. In contrast, dmPFC 4–7 Hz power was dynamically reduced when wild-type mice exhibited social approach behavior compared with leaving behavior with opposite moving directions. Replicating the social interaction-related LFP changes by oscillation-like optogenetic stimulation of PV interneurons was sufficient to increase social interaction in socially defeated mice.

Generally, studies utilizing an SI test have quantified the duration during which mice stayed in an IZ (Golden *et al*., 2011; Venzala *et al*., 2012; Ramaker & Dulawa, 2017). Our analysis failed to detect pronounced differences in LFP power between periods when mice stayed within and outside the IZ. These results were because animals’ behavioral patterns were not homogeneous across time in the IZ, suggesting a need to identify detailed behavioral patterns on a moment-to-moment basis to more accurately evaluate social interaction-related cortical LFP signals. Thus, we specifically extracted social approach behavior that more precisely represents animals’ psychological states with increased motivation and/or decreased anxiety than simple time spent in the IZ. Based on this identification of a behavioral pattern, we revealed pronounced reductions in 4–7 Hz power associated with social interactions in the dmPFC-BLA circuit.

Our results showed that dmPFC 4–7 Hz power increases occur during a target session, possibly representing a condition with increased attention against a target mouse, and the further increases occur during social avoidance in socially deficient mice. These results suggest that mice are inherently nervous and/or anxious for social interaction in a novel test environment and the strength of these mental states correlates with increased dmPFC 4–7 Hz power. This idea may partly be related to the previous observations that theta (4–10 Hz) power increases in the mPFC-BLA-ventral hippocampal circuit occurs in anxiogenic environments (Adhikari *et al*., 2010; Likhtik *et al*., 2014; Padilla-Coreano *et al*., 2019). In addition, recent studies have demonstrated that increases in mPFC-BLA 4 Hz power occur during fear retrieval (Dejean *et al*., 2016; Karalis *et al*., 2016; Caliskan & Stork, 2019; Ozawa *et al*., 2020), a condition that likely includes increased anxiety. Further studies are needed to determine whether these fear retrieval-related LFP signals are similar to the mPFC 4–7 Hz signals observed in this study. In addition, our results demonstrated that social interaction behavior is crucially associated with decreased dmPFC 4–7 Hz power, suggesting that the inherently increased dmPFC 4–7 Hz signals in anxiogenic environments need to be properly reduced to trigger social interaction behavior. This idea may be also related to the observations of decreased theta power in an open (less anxiogenic) environment (Dejean *et al*., 2016; Karalis *et al*., 2016; Caliskan & Stork, 2019; Ozawa *et al*., 2020). Indeed, a common contextual feature when taking social approach behavior observed in this study and passing into anxiogenic environments observed in the previous studies is that they both represent a context where mice need to overcome anxiety in relation to an open context and/or have motivation for exploration. The decreased dmPFC 4–7 Hz power might thus be partly explained by decreased anxiety.

Shank3 KO mice and socially defeated mice are utilized as an autism-like and a depression-like mouse model, respectively. A common behavioral symptom reported from these mouse models is decreased sociality, while they exhibit distinct behavioral features such as grooming and repetitive behaviors inherent in Shank3 KO mice (Peca *et al*., 2011; Mei *et al*., 2016). Our study demonstrated that both mouse models commonly exhibit larger increases in dmPFC 4–7 Hz power especially during social avoidance in a target session. According to the idea stated above, these pathological results are likely explained by continuous increased anxiety with the failure of dmPFC 4–7 Hz power reductions that prevent appropriate social interaction behavior. Taken together, our results suggest that reduced social interaction under pathological conditions is modulated by common neurophysiological mechanisms, increased and decreased dmPFC 4–7 Hz power during social avoidance and interactions, respectively.

A previous report has demonstrated that PFC 2–7-Hz oscillations entrain coherent activity between the amygdala and ventral tegmental area at the beta band and power changes in these oscillations during stress experiences predicts subsequent depression-like behavior (Kumar *et al*., 2014; Hultman *et al*., 2016). Consistent with their observation that normalization of these interregional oscillations by chemogenetic activation of prefrontal-amygdalar circuit reverses stress-induced behavioral deficits (Hultman *et al*., 2016), our results demonstrated that dynamic changes in dmPFC-BLA oscillations at the similar frequency band during behavior indeed correlate with social behavior and exogeneous inductions of such oscillations by optogenetic manipulations temporally induced social interaction behavior. An interesting remaining question is how these dynamically changing oscillations are linked with oscillations and coherent activity found in the other brain regions related to stress vulnerability and susceptibility, termed electome factors (Hultman *et al*., 2018).

We showed that dmPFC interneurons exhibited spike patterns phase-locked to 4–7 Hz oscillations. In addition, previous studies have demonstrated that PV interneurons are a crucial cell type for the generation of cortical gamma-range (30–60 Hz) oscillations (Whittington *et al*., 1995; Bartos *et al*., 2007; Cardin *et al*., 2009; Sohal *et al*., 2009; Buzsaki & Wang, 2012; Nakamura *et al*., 2015). These results suggest that inhibitory neurons in the dmPFC may be a main circuit for triggering social behavior-related LFP oscillations. In line with these ideas, we confirmed that 40-Hz photostimulation selective to PV interneurons could induce the same LFP power changes as those observed during social approach behavior, and this photostimulation led to a substantial increase in social interaction. While precise mechanisms for the reciprocal power changes in 4–7 Hz and 30– 60 Hz power by 40-Hz photostimulation remain unknown, this stimulus pattern seems effective in interfering with PV interneuronal spikes entrained by a dmPFC 4–7 Hz oscillation, leading to a reduction in 4–7 Hz power. Here, we note that as not all interneurons in the dmPFC are PV-positive interneurons and we did not directly demonstrate that the phase-locked interneurons expressed PV. Nonetheless, we at least demonstrated that mimicking these oscillatory patterns by utilizing the manipulation of PV-positive interneurons is sufficient to induce social interaction behavior that was reduced in defeated mice.

Accumulating evidence suggests that disruptions in E/I balances are crucial factors contributing to social behavior deficits in autism-like mouse models (Rubenstein & Merzenich, 2003; Helmeke *et al*., 2008; Yizhar *et al*., 2011; Selimbeyoglu *et al*., 2017). From the aspect of neuronal networks, our results add to a growing body of evidence demonstrating that dynamic changes in LFP oscillatory power regulated by inhibitory neuronal networks at appropriate timings are crucial for social behavior. The neurophysiological signatures found in our study may be helpful for a unified mechanistic understanding of the cellular- and network-based mechanisms underlying sociality and for identifying ASD-related and stress-induced pathophysiology that may lead to the amelioration of social behavior deficits.

## Materials and Methods

### Animals

All experiments were performed with the approval of the Experimental Ethics Committee at the University of Tokyo (approval number: P29-7 and P29-14) and according to the NIH guidelines for the care and use of mice.

Male C57BL/6J wild-type mice (8–10 weeks old) with preoperative weights of 20–30 g were used in this study. All the wild-type mice were purchased from SLC (Shizuoka, Japan). Shank3 knockout (KO) mice were provided by Shionogi & Co., Ltd. and 7 Shank3 KO mice were used for electrophysiological recordings at 8–10 weeks old. PV-IRES-Cre (PV-Cre) mice were obtained from Jackson Laboratory (B6;129P2-Pvalbtm1(cre)Arbr/J) and a subset of PV-Cre mice were subject to social defeat stress at 8–10 weeks old and then used for electrophysiological recordings. The animals were housed and maintained on a 12-h light/12-h dark schedule with lights off at 7:00 AM.

The Shank3 knockout mouse was designed with reference to the report by Peça et al (2011) with slight modifications. Briefly, a zinc-finger nuclease (ZFN) mRNA (Merck) was microinjected into the pronucleus of fertilized eggs of C57BL/6JJcl mice. The ZFN targets the following sequence of exon13 in the Shank3 gene; TGCTCCCCGCAGAAACcagagaGGACCGGACGAAGCG. The Shank3 deficient founder mice harboring 10 bases deletion in exon13 were identified by genome sequencing and inbreeded to produce homozygous deficient mice.

### Social defeat

Mice were exposed to chronic social defeat stress as previously described (Golden et al., 2011; Venzala et al., 2012; Ramaker & Dulawa, 2017; Abe et al., 2019). At least 1 week before beginning the social defeat experiment, all resident CD-1 mice (SLC, Shizuoka, Japan) more than 13 weeks of age were singly housed on one side of a home cage (termed the “resident area”; 42.5 cm × 26.6 cm × 15.5 cm). The cage was divided into two identical halves by a transparent Plexiglas partition (0.5 cm × 41.8 cm × 16.5 cm) with perforated holes, each with a diameter of 10 mm. The bedding in the resident area was left unchanged during the preoperative period. First, resident CD-1 mice were screened for social defeat experiments by introducing an intruder C57/BL6J mouse that was specifically used for screening into the home cage during three 3-min sessions on 3 subsequent days. Each session included a different intruder mouse. CD-1 mice were selected as aggressors in subsequent experiments based on three criteria: during the three 3-min sessions, (1) the mouse attacked in at least two consecutive sessions, (2) the latency to initial aggression was less than 60 s, and (3) the above two criteria were met for at least two consecutive days out of three test days. After screening, an experimental intruder mouse was exposed to social defeat stress by introducing it into the resident area for a 7–10-min interaction. The interaction period was immediately terminated if the intruder mouse had a wound and bleeding due to the attack, resulting in interaction periods of 7–10-min. After the physical contact, the intruder mouse was transferred across the partition and placed in the opposite compartment of the second resident home cage for the following 24 h; this allowed the intruder mouse to have sensory contact with the resident mouse without physical contact. Over the following 10-day period, the intruder mouse was exposed to a new resident mouse so that the animals did not habituate the same residents.

### Surgical procedures

A single surgery was performed in each animal for either (i) implantation of a tetrode assembly or (ii) implantation of a tetrode assembly and optic fibers followed by injection of a virus vector. During all surgeries, the animals were anesthetized with isoflurane gas (1–3%), and circular craniotomies were made using a high-speed drill at the indicated coordinates. (i) For LFP recordings without spikes, an electrode assembly that consisted of 3 and 4 immobile tetrodes was stereotaxically implanted above the dmPFC (2.00 mm anterior and 0.50 mm lateral to bregma) at a depth of 1.40 mm and the BLA (0.80 mm posterior and 3.00 mm lateral to bregma) at a depth of 4.40 mm, respectively. The tetrodes were constructed from 17-μm-wide polyimide-coated platinum-iridium (90/10%) wires (California Fine Wire), and the electrode tips were plated with platinum to lower the electrode impedances to 200–250 kΩ. Stainless steel screws were implanted on the skull and attached to the cerebellar surface to serve as ground/reference electrodes. For spike recordings, an electrode assembly that consisted of 6 independently movable tetrodes was stereotaxically implanted above the mPFC (1.94 mm anterior and 0.83 mm lateral to bregma). (ii) For optogenetic experiments, 300 nl AAV5-EF1a-DIO-ChR2-YFP (UNC Vector Core, 1.0 × 1013 vg/ml) was injected into the dmPFC (2.00 mm anterior and 0.50 mm lateral to bregma at a depth of 1.40 mm) over 3 min in a PV-Cre mouse, and an optical fiber (core diameter = 200 µm) was then implanted into the same region. In addition, an electrode assembly for LFP recordings was implanted as described above. Finally, all devices were secured to the skull using stainless steel screws and dental cement. After all surgical procedures were completed, anesthesia was discontinued, and the animals were allowed to spontaneously awaken. Following surgery, each animal was housed in a transparent Plexiglas cage with free access to water and food for more than one week.

For spike recordings, the tetrodes were advanced to the targeted brain regions over a period of at least one week following surgery. The depth of the electrodes was adjusted while the mouse rested in a pot placed on a pedestal. The electrode tips were advanced up to 62.5 μm per day over a period of at least 10 days following surgery. The tetrodes were then settled into the targeted area so that stable recordings were obtained.

### Electrophysiological recording

The mouse was connected to the recording equipment via Cereplex M (Blackrock), a digitally programmable amplifier, which was placed close to the animal’s head. The output of the headstage was conducted to the Cereplex Direct recording system (Blackrock), a data acquisition system, via a lightweight multiwire tether and a commutator. For recording electrophysiological signals, the electrical interface board of the tetrode assembly was connected to a Cereplex M digital headstage (Blackrock Microsystems), and the digitized signals were transferred to a Cereplex Direct data acquisition system (Blackrock Microsystems). Electrical signals were sampled at 2 kHz and low-pass filtered at 500 Hz. The unit activity was amplified and bandpass filtered at 750 Hz to 6 kHz. Spike waveforms above a trigger threshold (50 μV) were time-stamped and recorded at 30 kHz in a time window of 1.6 ms. The animal’s moment-to-moment position was tracked at 15 Hz using a video camera attached to the ceiling. The frame rate of the movie was downsampled to 3 Hz, and the instantaneous speed of each frame was calculated based on the distance traveled within a frame (∼333 ms). In the following analyses, video frames with massive optical noise or periods that were not precisely recorded due to temporal breaks of image data processing were excluded. All recordings from a behavioral task were performed once so that all the tasks were novel for the mice and no duplications of samples were thus included.

### Social interaction (SI) test

Social interaction tests were performed inside a dark room with a light intensity of 10 lux in a square-shaped box (39.3 cm × 39.3 cm) enclosed by walls 27 cm in height. A wire-mesh cage (6.5 cm × 10 cm × 24 cm) was centered against one wall of the arena during all social interaction sessions. Each social interaction test included two 150-s sessions (separated by an intersession interval of 30 s) without and with the target CD-1 mouse present in the mesh cage; these sessions were termed no target and target sessions, respectively. In the no-target session, a test C57BL/6J mouse was placed in the box and allowed to freely explore the environment. The C57BL/6J mouse was then removed from the box. In the 30-s break between sessions, the target CD-1 mouse was introduced into the mesh cage. The design of the cage allowed the animal to fit its snout and paws through the mesh cage but not to escape from the cage. In the target session, the same C57BL/6J mouse was placed beside the wall opposite to the mesh cage. For optogenetic experiments, the duration of a target session was 10 min. In each session, the time spent in the interaction zone (IZ), a 14.5 cm × 24 cm rectangular area extending 8 cm around the mesh cage. The social interaction (SI) ratio was computed as the ratio of time spent in the interaction zone in the target session to the time spent there in the no-target session. Social avoidance zones are defined as 9.0 cm × 9.0 cm square areas projecting from both corner joints opposing the cage.

### Open field (PF) test

Mice were placed in an open field (39.3 cm × 39.3 cm) with a wall height of 50 cm and allowed to move freely throughout the field for 10 min. The floor was illuminated with an overhead light, which produced light intensities of 25 lux in the field. A border area and center area were defined as areas less than and more than 8 cm from the wall, respectively.

### Optogenetics

The mice underwent one of the following photostimulation protocols during a target session for 10 min: (1) 40-Hz stimulation with 12.5-ms blue light pulses (472 nm, ∼3 mW output from fiber) at 40 Hz applied with a periodicity of 30 s, (2) 10-Hz stimulation with 50-ms blue light pulses at 10 Hz applied with a periodicity of 30 s, and (3) 4-Hz stimulation with 125-ms blue light pulses at 4 Hz applied with a periodicity of 30 s. In a representative example shown in Figure 6B and the analysis in Figure 6C and 6D, each photostimulation was applied for 10 s.

For behavioral experiments, a test day for each mouse includes a sequence of a target session with no photostimulation (no stim), and target sessions with 4-Hz, 10-Hz, and 40-Hz photostimulation. The order of these sessions was similar in all mice. Each session lasted for 10 min, during which photostimulation was applied for 2 s every 30 s, and overall changes in the animal’s social interactions throughout the session were examined.

### Histological analysis to confirm electrode placement or cannula placement

The mice were overdosed with isoflurane, perfused intracardially with 4% paraformaldehyde in phosphate-buffered saline (pH 7.4) and decapitated. After dissection, the brains were fixed overnight in 4% PFA and equilibrated with 20% and 30% sucrose in phosphate-buffered saline for an overnight each. Frozen coronal sections (100 μm) were cut using a microtome, and serial sections were mounted and processed for cresyl violet staining. For cresyl violet staining, the slices were rinsed in water, stained with cresyl violet, and coverslipped with Permount. The positions of all electrodes were confirmed by identifying the corresponding electrode tracks in histological tissue.

### LFP analysis

To compute the time-frequency representation of LFP power, LFP signals were convolved using complex Morlet wavelet transformation at frequencies ranging from 1 to 250 Hz. The absolute power spectrum of the LFP during each 10-ms time window was calculated, and z-scores were computed for each frequency band across an entire period including the no-target and target sessions. In Figures 2 and 3, the ratio of absolute power during a target session to that during a no-target session at a specific frequency band (4–7 Hz or 30–60 Hz) was computed. When data were obtained from multiple electrodes in a mouse, they were averaged to single values within each mouse.

### Definition of social approach and leaving behavior

For each location on the animal’s trajectories (bin = 1 s), a cage-oriented moving direction θ was computed as an angle between the animal’s moving direction at the location and a straight line connecting the location of the animal and the center of the cage. Angles θ = 0° and 180° indicate moving directly toward and away from the center of the cage, respectively. Candidate periods of social approach and leaving behavior were defined when the animals stayed in the half of the field containing the IZ and the cage-oriented moving directions were less than and more than 90°, respectively. If a candidate period lasted for more than 5 s, the first 5-s period was regarded as a social approach and leaving behavior. The periods that were not classified as social approach and leaving behavior were termed the other periods.

### Spike unit analysis

Neurons recorded from all electrodes targeting the dmPFC were included in this analysis. Spike sorting was performed offline using the graphical cluster-cutting software Mclust. Sleep recordings obtained before and after the behavioral paradigms were executed were included in the analysis to assure recording stability throughout the experiment and to identify cells that were silent during behavior. Clustering was performed manually in 2D projections of the multidimensional parameter space (i.e., comparisons between the waveform amplitudes, the peak-to-trough amplitude differences, the waveform energies, and the first principal component coefficient (PC1) of the energy-normalized waveform, each measured on the four channels of each tetrode). Only clusters that could be stably tracked across all behavioral sessions were considered to be the same cells and were included in our analysis. Similar to classification criteria in the PFC reported in previous studies (Wilson *et al*., 1994; Tierney *et al*., 2004; Homayoun & Moghaddam, 2007), neurons with baseline spike rates of >10 Hz were classified as putative fast-spiking (FS) interneurons, whereas neurons with baseline spike rates of <10 Hz and spike width of >0.6 ms were classified as putative regular-spiking (RS) pyramidal neurons. No *a priori* power analyses were performed to determine sample sizes. Experiments were instead designed to encompass a comparable number of cells as several previous studies of spike-phase computation among prefrontal principal cells (e.g. (Karalis *et al*., 2016; Abe *et al*., 2019; Okonogi & Sasaki, 2021)).

For each cell, the degree of phase locking during a target session was analyzed. For approach or leaving behavior, the phase-spike rate distribution was computed by plotting the firing rate as a function of the phase of 1–4 Hz, 4–7 Hz, 7–15 Hz, 15–30 Hz, and 30–60 Hz LFP traces, divided into bins of 30° and smoothed with a Gaussian kernel filer with standard deviation of one bin (30°), and a Rayleigh r-value was calculated as mean vector length (MVL).

### Statistical analysis

All data are presented as the mean ± standard error of the mean (SEM), unless otherwise specified, and were analyzed using Python and MATLAB. For normally-distributed data, individual data points are displayed in addition to sample mean and standard errors of the mean or presented in the sourcedata.xlsx. For non-normally-distributed data, data are displayed as distributions, with data points presented in the sourcedata.xlsx. For each statistical test, data normality was first determined by the F test, and non-parametric tests applied where appropriate. Comparisons of two-sample data were analyzed by paired t-test and Mann-Whitney U test. Multiple group comparisons were performed by post hoc Bonferroni corrections. The null hypothesis was rejected at the *P* < 0.05 level.

### Data availability

All data generated or analysed during this study are included in the manuscript and supporting file; Source Data files have been provided for main and supplementary Figures. Original datasets are available at https://www.dropbox.com/s/66fi2hgayj2nh5v/Kuga%20et%20al%202021%20datasets-20211102T181358Z-002.zip?dl=0

## ACKNOWLEDGMENTS

This work was supported by KAKENHI (19H04897; 20H03545) from the Japan Society for the Promotion of Science (JSPS), a grant (JPMJCR21P1) from the JST CREST, and a grant (1041630; JP21zf0127004) from the Japan Agency for Medical Research and Development (AMED) to T. Sasaki; funds from the JST Exploratory Research for Advanced Technology (JPMJER1801), and Institute for AI and Beyond of the University of Tokyo to Y. Ikegaya.

## AUTHOR CONTRIBUTIONS

N.K., R.A. and T.S. designed the study. N.K., R.A., K.T., and T.S. acquired the electrophysiological data. N.K., R.A., and T.S. performed the analyses. Y.I. supervised the project. T.S. prepared the figures and wrote the main manuscript text, and all the authors reviewed the main manuscript text.

## DECLARATION OF INTERESTS

The authors declare no competing interests.

---

**Figure S1.**
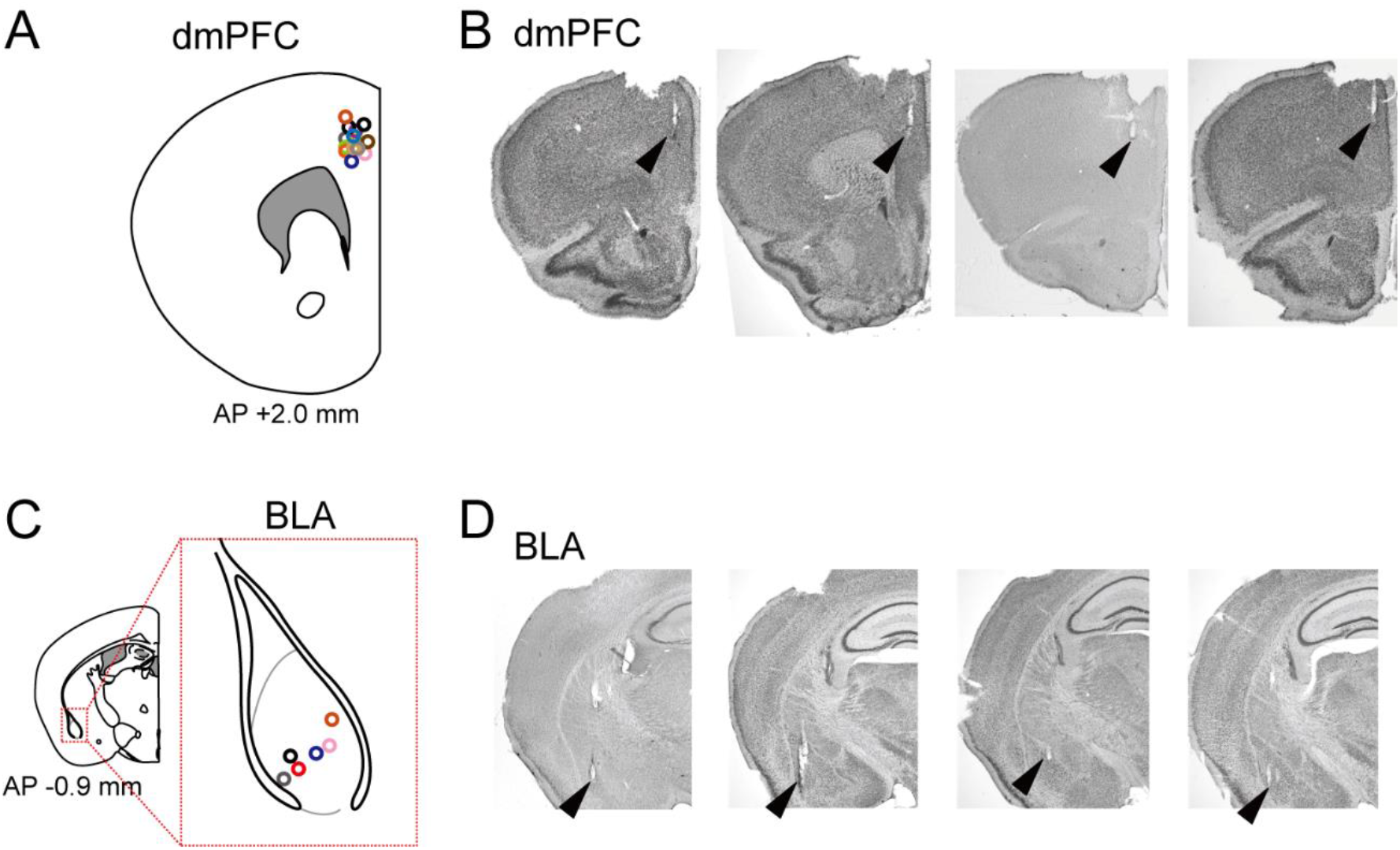
Confirmation of recording sites. (A) Superimpositions of recording sites for all electrodes (from 14 mice shown in Figure 1) indicated by circles on the dmPFC. (B) Typical pictures of cresyl violet-stained sections with the arrowheads indicating the tips of electrode tracks. (C) Same as A but for the BLA (from 6 mice shown in Figure 1). The gray dotted line represents the border to define the BLA region. (D)) Same as B but for the BLA.

**Figure S2.**
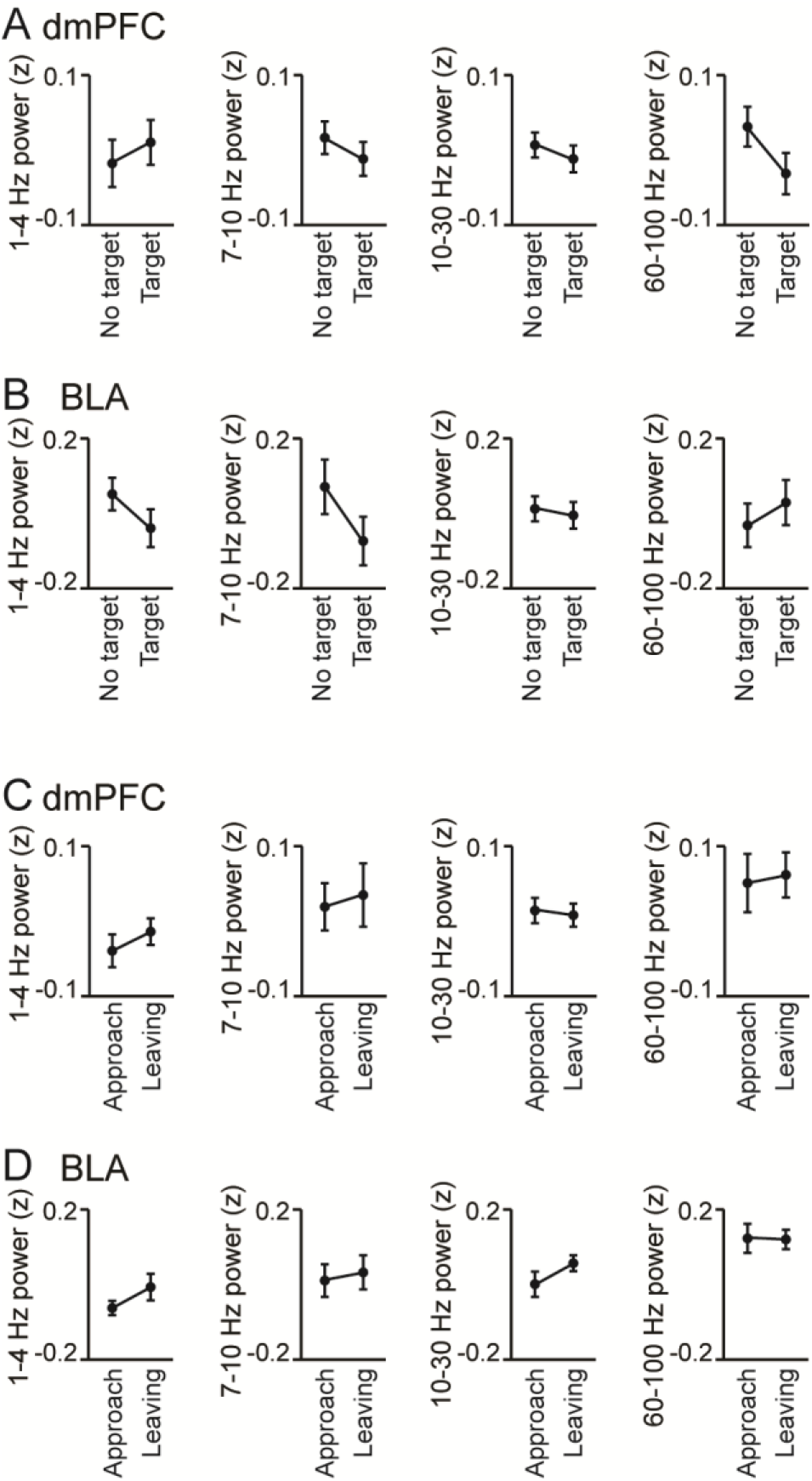
Analyses for the frequency bands other than 4–7 Hz and 30–60 Hz. (A, B) Supplementary datasets for Figure 1F and 1G. For the frequency bands of 1–4 Hz, 7– 10 Hz, 10–30 Hz, and 60–100 Hz, LFP power in the dmPFC (A) and BLA (B) was compared between the target and no target sessions (*n* = 14 and 6 mice, respectively). Data are presented as the mean ± SEM. No significant differences were found in these comparisons (*P* > 0.05, paired *t*-test, taget vs no target). (C, D) Supplementary datasets for Figure 4F and 4G. For the frequency bands of 1–4 Hz, 7– 10 Hz, 10–30 Hz, and 60–100 Hz, LFP power in the dmPFC (C) and BLA (D) was compared power between approach and leaving behavior (*n* = 14 and 6 mice respectively). Data are presented as the mean ± SEM. No significant differences were found in these comparisons (*P* > 0.05, paired *t*-test, approach vs leaving).

**Figure S3.**
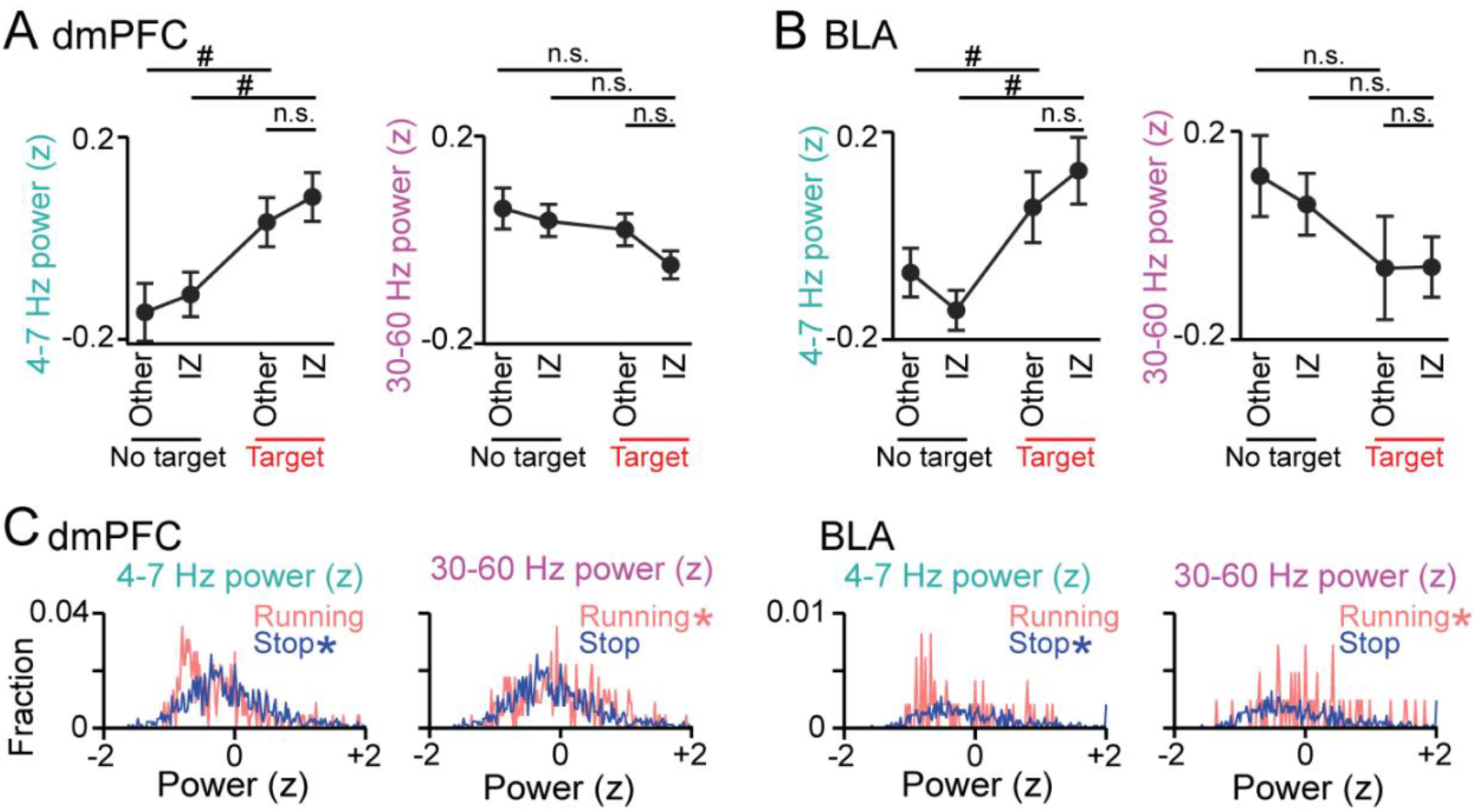
LFP power changes in the target session. (A, B) No significant changes in dmPFC and BLA LFP power between the IZ and the other areas in the target session. In the dmPFC and BLA, 4–7 Hz (left) and 30–60 Hz (right) power was compared between the IZ and the other areas (A; dmPFC, *n* = 14 mice; 4–7 Hz: *t*_13_ = 0.76, *P* = 0.46; 30–60 Hz: *t*_13_ = 1.38, *P* = 0.19; B; BLA, *n* = 6 mice; 4–7 Hz: *t*_5_ = 0.76, *P* = 0.48; 30–60 Hz: *t*_5_ = 0.02, *P* = 0.98). Data are presented as the mean ± SEM. #*P* < 0.05, paired *t*-test followed by Bonferroni correction. (C) Comparisons of distributions of LFP power between running (>5 cm/s) and stop (<1 cm/s) periods in the target session (dmPFC, *n* = 254 and 1008; 4–7 Hz: *Z* = 4.93, *P* = 8.26 × 10^-7^; 30–60 Hz: *Z* = 2.13, *P* = 0.033; BLA, *n* = 58 and 519; 4–7 Hz: *Z* = 2.42, *P* = 0.016; 30– 60 Hz: *Z* = 2.00, *P* = 0.045, Mann-Whitney U test). Asterisks represent significantly higher power. (D)) Same as B but for the BLA.

**Figure S4.**
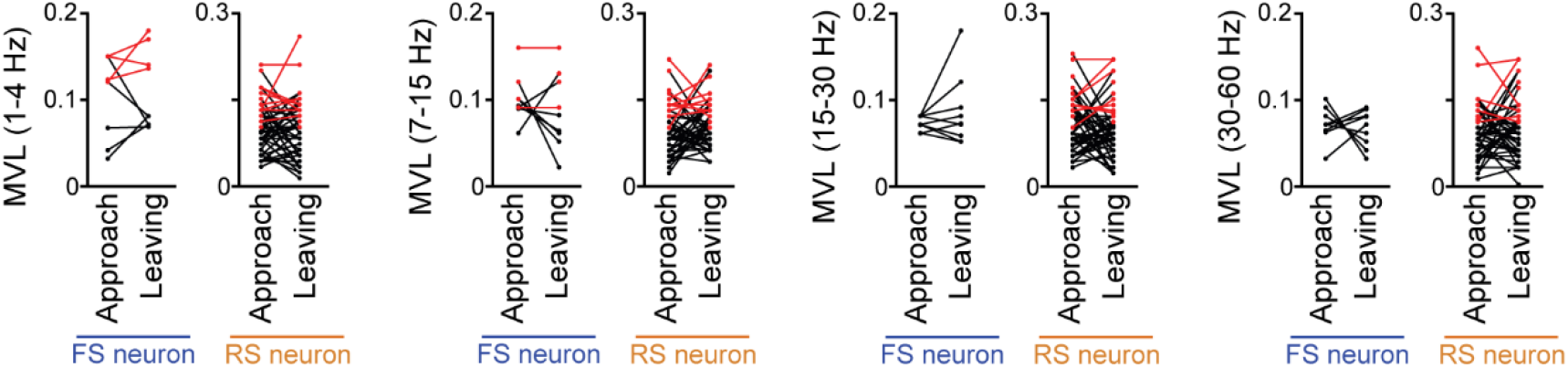
Comparisons of MVL for the oscillations (at 1–4 Hz, 4–7 Hz, 7–15 Hz, 15–30 Hz, and 30–60 Hz) other than the 4–7-Hz LFP oscillations between approach and leaving behavior. Data are shown same as in Figure 5C. RS and FS neurons were separately analyzed. Each dot and line represent each neuron. The red dots indicate significant MVL, computed from shuffled datasets. The red lines indicate neurons showing significant MVL in both periods, which were considered as behavior-irrelevant phase-locked neurons and excluded from statistical analyses. In all comparisons, no statistical differences were observed (*P* > 0.05) (1–4 Hz: FS: *n* = 5 neurons; *t*_4_ = 0.33, *P* = 0.76; RS: *n* = 39 neurons; *t*_38_ = 0.28, *P* = 0.78; 7–15 Hz: FS: *n* = 7 neurons; *t*_6_ = 0.86, *P* = 0.42; RS: *n* = 39 neurons; *t*_38_ = 1.16, *P* = 0.25; 15–30 Hz: FS: *n* = 9 neurons; *t*_8_ = 1.06, *P* = 0.32; RS: *n* = 41 neurons; *t*_40_ = 1.69, *P* = 0.18; 30–60 Hz: FS: *n* = 9 neurons; *t*_8_ = 0.42, *P* = 0.69; RS: *n* = 44 neurons; *t*_43_ = 0.64, *P* = 0.52; paired *t*-test).

## Notes

### Competing Interest Statement

The authors have declared no competing interest.

